# Comprehensive Assessment of Sequence and Expression of Circular-RNAs in Progenitor Cell Types of the Developing Mammalian Cortex

**DOI:** 10.1101/553495

**Authors:** Martina Dori, Leila Haj Abdullah Alieh, Daniel Cavalli, Simone Massalini, Mathias Lesche, Andreas Dahl, Federico Calegari

**Affiliations:** DFG-Research Center and Cluster of Excellence for Regenerative Therapies (CRTD), School of Medicine, Technische Universität Dresden; Fetcherstrasse 105, 01307, Dresden, Germany; Center for Genome Research, Department of Life Sciences, University of Modena and Reggio Emilia; Via G. Campi, 287, 41100, Modena, Italy; Center for Molecular and Cellular Bioengeneering (CMCB), DFG-Research Center for Regenerative Therapies (CRTD), Technische Universität Dresden; Fetcherstrasse 105, 01307, Dresden, Germany

## Abstract

Circular (circ) RNAs have recently emerged as a novel class of non coding transcripts whose identification and function remain elusive. Among many tissues and species, the mammalian brain is the organ in which circRNAs are more abundant and first evidence of their functional significance started to emerge. Yet, even within this well studied organ, annotation of circRNAs remains fragmentary, their sequence is unknown and their expression in specific cell types was never investigated. Overcoming these limitations, here we provide the fist comprehensive identification of circRNAs and their expression patterns in proliferating neural stem cells, neurogenic progenitors and newborn neurons of the developing mouse cortex. Extending the current knowledge about the diversity of this class of transcripts by the identification of nearly 4,000 new circRNAs, our study is the first to provide the full sequence information and expression patterns of circRNAs in cell types representing the lineage of neurogenic commitment. We further exploited our data by evaluating the coding potential, evolutionary conservation and biogenesis of circRNAs that we found to arise from a specific sub-class of linear mRNAs. Our study provides the arising field of circRNA biology with a powerful new resource to address the complexity and potential biological significance of this new class of transcripts.

## INTRODUCTION

In the last few decades, the field of RNA biology has witnessed impressive developments. Fuelled by new sequencing technologies, these included the comprehensive annotation of micro and long non-coding (lnc) RNAs in various organisms and tissues, the characterization of RNA modifications and the new field of *epitranscriptomic* and the discovery of an entirely new class of non coding RNAs: circular (circ) RNAs ^1^.

CircRNAs are transcripts whose 3’ and 5’ ends are covalently linked in a non-linear manner resulting in a so-called backsplice junction ^2,3^. The lack of a 3’ poly(A) tail and 5’ capping provides this class of RNAs resistance to exonuclease activity and, thus, an average longer half-live as compared to linear RNAs ^4–6^. Transcripts with these characteristics have long been known but until recently circRNAs were primarily found in viruses ^7^ and although some reports indicated their origin also from eukaryotic genomes ^8–10^ these were still considered a rarity or a byproduct of splicing with no specific function. This view was completely changed very recently after the identification of thousands of circRNAs including some with regulatory functions during brain development ^6,11–14^.

Despite their abundance, predicting circRNAs remains burdensome and typically relies on bioinformatic tools identifying sequences across backsplice junctions from RNA sequencing data obtained upon depletion of ribosomal RNA ^15^. Although this has resulted in the prediction of thousands of potential circRNAs in cell lines or whole organs of many species ^6,11,13,16–19^, this approach based on ribosomal RNA depletion has the critical limitation that reads that do not map on a backsplice junction cannot be assigned to a circular, as opposed to a linear, transcript resulting in the exclusion of the overwhelming majority of the sequencing data. In turn, this makes it impossible to reliably reconstruct neither the full sequence nor the expression level of large pools of circRNAs. Overcoming these limitations, the use of the exonuclease RNase R triggers the digestion of linear RNAs thereby allowing the isolation of circRNAs ^4^. To date, this strategy was applied to a few cell lines or whole organs ^6,16,20–22^ but the expression patterns of circRNAs in specific cell types in physiological conditions is not known and a comprehensive reconstruction of their sequence is yet to be reported.

In addition to a poor classification, barely a handful of circRNAs have been suggested to be functionally relevant. For example, and beside their use as biomarkers in various diseases from cancer to diabetes ^23–26^, a study concluded that at least some circRNA, such as circZNF609, may retain coding potential ^27^. Additionally, Drosophila’s circMbl was found to interact with MBL protein resulting from the linear form of the same transcript ^20^. Finally, and perhaps as the only circRNA-mediated molecular mechanism underlying a specific cellular effect, Cdr1as/ciRS-7 was shown to act as a sponge for miR-7 during development ^12,13^ and its depletion altered synaptic transmission in adulthood ^14^.

From these and other studies, it emerged that the mammalian brain is the organ most enriched in circRNAs ^28,29^. Hence, given the scarce knowledge about circRNA function and lack of studies reporting their sequence and expression in specific cell types, we here decided to exploit a double reporter mouse line previously characterized by our group and allowing the isolation of proliferating neural stem cells, neurogenic progenitors and newborn neurons based on the combinatorial expression of RFP and GFP, respectively ^30^.

Specifically, during cortical development, proliferative progenitors (PP) progressively switch from divisions that expand their population to divisions that generate more committed, differentiative progenitors (DP), which in turn are consumed to generate newborn neurons (N) ^31,32^. Hence, to identify the three coexisting subpopulations of PP, DP and N during mouse corticogenesis, our group has generated a double reporter mouse line expressing i) RFP under the control of the *Btg2* promoter and identifying the switch of PP to DP and ii) GFP under the control of the *Tubb3* promoter as a marker of newborn N ^30^. Validating this approach, the combinatorial expression of the two reporters allowed the isolation of PP (RFP−/GFP−) from DP (RFP+/GFP−) and N (GFP+, irrespective of RFP) and identification and validation of transcription factors and lncRNAs functionally involved in neurogenic commitment ^30,33,34^. Hence, we decided to further exploit this mouse line and provide the arising field of circRNA biology with the first resource identifying circRNAs sequence and expression patterns in specific cell types of the developing mammalian cortex.

## RESULTS

### Comprehensive identification of cell type-specific circRNAs of the mouse cortex

To identify cell type-specific circRNAs we FAC-sorted PP, DP and N, each in three biological replicates, from the mouse cortex at embryonic day (E) 14.5 as previously described ^30^. Total RNA was then treated with RNase R to degrade linear RNAs, which we found to be very efficient in reducing the levels of linear transcript down to undetectable levels even among those with the highest expression levels and most stable predicted secondary structure (**Figure S1A**). This was then followed by 150 bp single-end, strand-specific, high-throughput sequencing. Reads were then aligned to the reference mouse genome (mm9) and unmapped reads used to identify the circularizing, backsplice junctions predictive of putative circRNAs (**Figure S1B**).

As a first step to assess the sequence of circRNAs and their expression in cortical cell types, we collected their genomic coordinates putting together all replicates of PP, DP and N and obtaining an initial set of 6,033 putative circRNAs (**Figure 1A**). Then, according to their genomic locations, we selected genic transcripts whose start and end sites coincided with the annotated start and end site of an exon and separated them from the remaining transcripts that included genic transcripts with ends not coinciding with exons, antisense or intergenic ones (**Figure S1D**). Within the former, we separated sequences belonging to introns from exons while the latter were considered as a single exon. Finally, we calculated the relative RPKM value of each intron and exon (**Figure 1A** and Materials and Methods).

**Figure 1.**
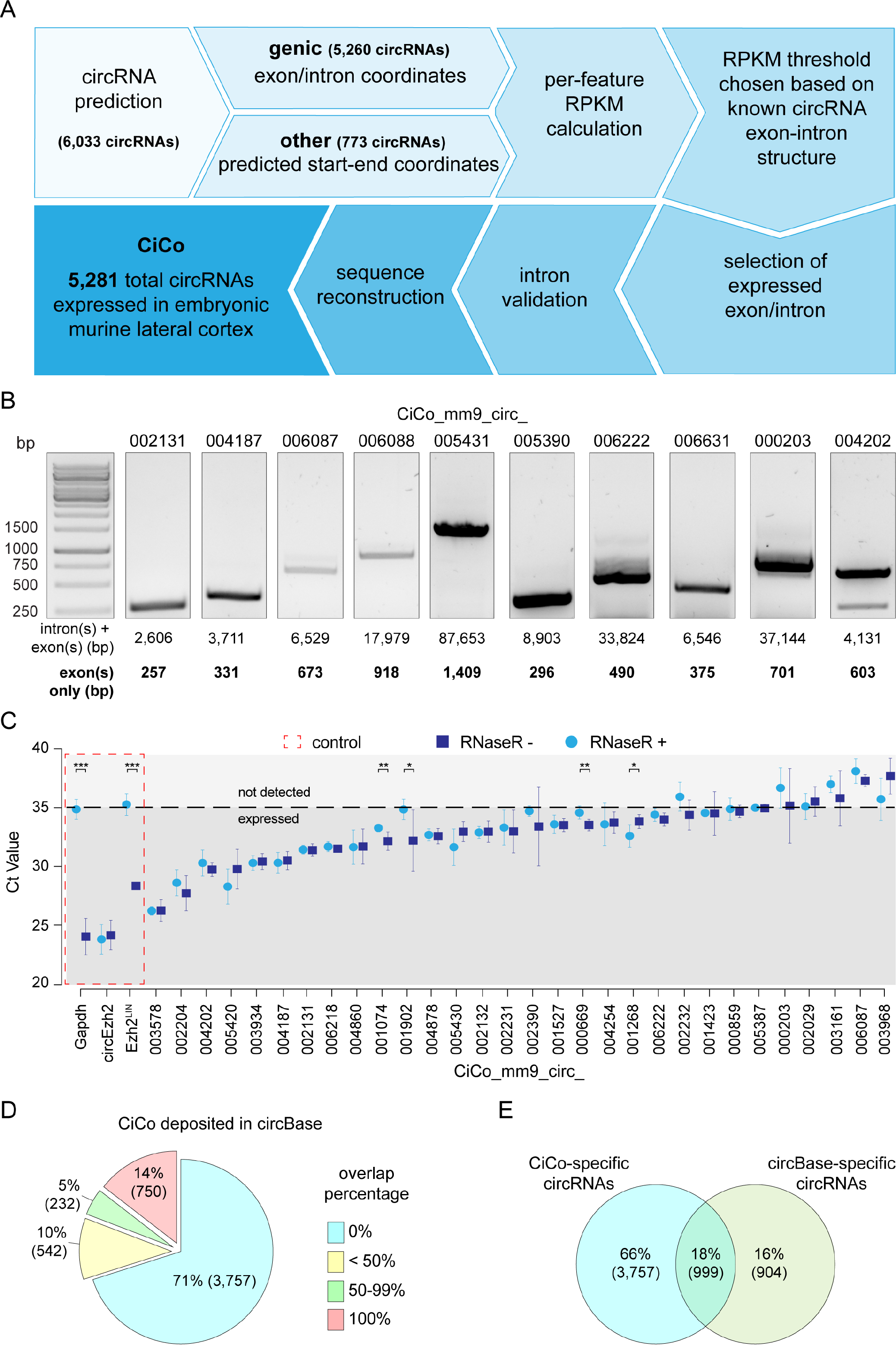
circRNA prediction, sequence determination and validation. (**A**) Strategy for the identification of circRNAs and determination of their sequence and expression in cortical cell types. (**B**) DNA gel electrophoresis upon PCR amplification of 10 circRNAs identified as in A. Full gels are available in **Supplementary Figure S2**. Note the detection of bands with a molecular weight consistent with the expected exonic, but not intronic, regions. (**C**) Graph representing the cycle threshold (Ct) values upon RT-PCR of total RNA from the E14.5 mouse cortex, with or without a prior treatment with RNase R (as indicated) and using divergent primers for 30 predicted circRNAs and linear and circular controls (red dashed line). Dashed black line indicates the Ct threshold of detection. N=2, n=3; bars=SDs (* *p*<0.05; ** *p*<0.01; *** *p*<0.001; Student’s t-test). (**D**) Distribution of CiCo’s entries overlapping circRNAs deposited in circBase and (**E**) Venn Diagram representing the proportion of circBase- and CiCo-specific circRNAs as well as common ones.

Given the need to establish an unbiased minimum threshold of RPKM to define “*expression*”, we next selected 10 predicted genic circRNAs from our dataset including 6 that were not described by previous studies. We then cloned and sequenced these circRNAs from RNase R-treated lysates from the E14.5 mouse lateral cortex and found that in all cases the predicted exon(s) (2-15 for each circRNA; 45 in total) were included in their sequence while not a single intron (35 in total) could be detected (**Figure 1B**). Hence, we chose the highest RPKM value previously calculated among these predicted, but not detected, introns as a minimum threshold to define expression (RPKM>5.5). As a result, this allowed us to redefine among the original list of 6,033 putative circRNAs the 5,281 fulfilling our criteria for expression (**File S1**). Furthermore, we selected form this list 6 genic circRNAs for which introns were predicted as part of their sequence and designed divergent primers to validate their existence. Again, while confirming the presence of exons in these circRNAs, we were unable to confirm intronic sequences (**Figure S1C**) leading us to conclude that our list of genic circRNAs are primarily exonic and that reads mapping on intronic locations may derive from lariats.

Next, as a validation of our approach, we selected 30 circRNAs among this list of 5,281 including 9 belonging to the bottom 30% and 4 to the bottom 10% in expression levels. Subsequent RT-PCRs were performed from the E14.5 mouse cortex, with or without RNase R treatment, using either divergent primers spanning over the backsplice junctions (**Figure S1E**) or convergent primers for 2 linear transcripts used as internal negative control of the RNase R treatment (GAPDH and Ezh2^LIN^). This confirmed the presence of the vast majority (80%) of the selected circRNAs (**Figure 1C**). Importantly, failure to detect some circRNAs was independent from their predicted expression levels pointing out more a suboptimal choice of primers than false positives in our analysis. From here, we then reconstructed the comprehensive list, sequence and expression levels of the 5,281 circRNAs detected in PP, DP and N and representing the neurogenic lineage during corticogenesis that we named **<u>CiCo</u>** for **<u>Ci</u>**rcRNAs of the mouse **<u>Co</u>**rtex (**Figure 1A** and **File S1)**.

We next compared CiCo with the most complete resource of circRNAs available: circBase ^35^. In particular, due to the stage- and tissue-specific expression of circRNAs, we selected from circBase murine circRNAs predicted from whole embryos and brains. Considering total or partial overlaps, down to the single nucleotide, as known circRNAs we found that our dataset extended the list of known circRNAs by nearly 3-fold adding to the ~1,900 transcripts of circBase ~3,700 new circRNAs of CiCo (**Figure 1D**) for a total of ~5,600 unique circRNAs (**Figure 1E**). As previously shown in the case of lncRNAs ^30,33^, this high rate of novel circRNAs found in our study relative to previous reports highlights the power of cell type-specific analyses allowing the identification of transcripts enriched in specific cell populations and being diluted out when considering bulk tissues or whole organs. In turn, this suggests a new layer of complexity in circRNA biology as these appeared to be not only stage- and tissue-specific, but also within the same developmental stage and tissue, to retain highly specific expression in individual cell types, which we sought to dissect next.

### General features and differential expression of CiCo

Previous studies reported that ~80% of the predicted circRNAs overlap in the sense strand with genes ^6,11,13^. In our dataset 96% of the expressed sequences overlapped genes with the remaining 4% including either antisense or intergenic circRNAs (**Figure 2A**). This seemed in contrast with the similar proportion of sense and antisense transcripts among lncRNAs ^36^, which can be explained by the overall lower expression of this class of transcripts biasing against the detection of circRNAs derived from them. Regarding length distribution and exon density, we found that most circRNAs (90%) were less than 1 kb long, primarily 250-500 bp, and including on average 2-3 exons (**Figure 2B**). As expected, a linear correlation was found between the length of circRNAs and the number of exons that they included with no specific bias in their distribution across or within chromosomes (data not shown).

**Figure 2.**
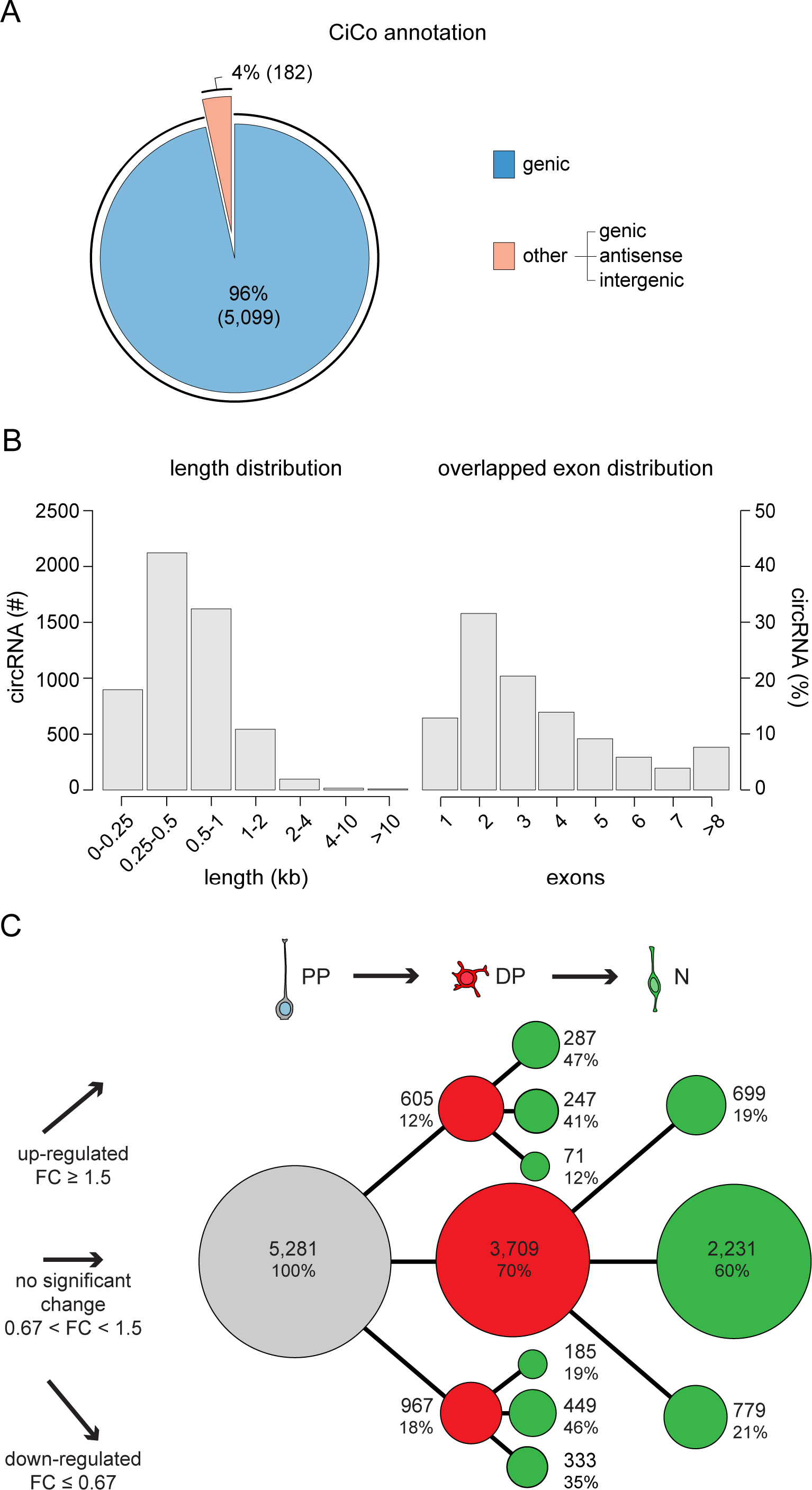
General features of CiCo. (**A**) Number and proportion of CiCo and their genomic features almost completely (96%) from genic regions. (**B**) Length (left) and exon number (for genic transcripts, right) distribution of circRNAs. (**C**) Differential expression (50% i.e.: FC ≥1.5 or ≤0.67; no FDR being applied) of CiCo expressed in PP (grey), DP (red) and N (green). Abundance and proportion of circRNAs detected in each cell type are indicated and represented proportionally to the area of circles and pattern of expression.

Next, we analysed CiCo’s expression profiles of PP, DP and N. In particular, we focused on the differentially expressed circRNAs considering a 50% threshold (i.e. a fold change (FC) by ≥1.5 or ≤0.67 for up- or down-regulation, respectively) (**Figure 2C**). While a substantial proportion (42%) of circRNAs showed no significant change in expression among cell types, subdividing differentially expressed circRNAs into the possible patterns of up- or down-regulation during the neurogenic lineage revealed a distribution that was remarkably similar to that found in linear transcripts ^30^ (**Figure 2C**). In addition, when analysing the biological functions and processes of the parental genes enriched in each group, we found that circRNAs up-regulated during neurogenesis were primarily associated to synaptogenesis and neuronal development. Conversely, down-regulated circRNAs were associated to cell cycle and regulation of transcription (data not shown).

Among transcripts showing specific expression patterns, an intriguing group of circRNAs emerged that was transiently up- or down-regulated specifically in DP compared to both PP and N (69 and 182, respectively) (**Figure 2C**). As previously shown by our group ^30^, these patterns of expression account for a small, underrepresented proportion of differentially expressed transcripts (1-2%). Yet, at least among mRNAs and lncRNAs, many from this small subset of genes revealed to play key roles in corticogenesis ^30,34^ raising the possibility that this also applies to circRNAs here identified for the first time.

### Properties of circRNAs: miRNA-sponging potential, translation and evolutionary conservation

Our novel assessment of the sequence of circRNAs gave us the possibility to gain new insights into their putative function(s). Since sponging of miRNAs by Cdr1as was the first suggested role for a circRNA ^12,13^, we searched within CiCo for miRNA seed sequences. When assessing the highest possible number of seed sequences present in any given circRNA that would potentially target any given miRNA, we found a very broad range of sponging properties reaching up 99 seeds in Cdr1as (CiCo ID 006933) for miR-7b and including one seed for miR-671, as previously described ^13^. Despite this extreme example, only a handful of circRNAs displayed a number of seeds for a single miRNA that exceeded 5 (Figure 3A). Moreover, since this analysis did not account for the potential of circRNAs to sponge more miRNAs independently from the number of seeds targeting each individual miRNA, we identified in CiCo ID 006344 the circRNA with the most predicted seeds for a total of nearly 1,400 miRNAs of which 4 (miR-1187, 466i, 466k and 669c) with a remarkably high number of seeds (46, 43, 37 and 30 respectively; **Figure 3A** and **File S1**). However, such a high promiscuity in miRNA binding sites and limitations in the bioinformatic tools to predict sponging properties makes it difficult to infer the biological significance of this circRNA.

**Figure 3.**
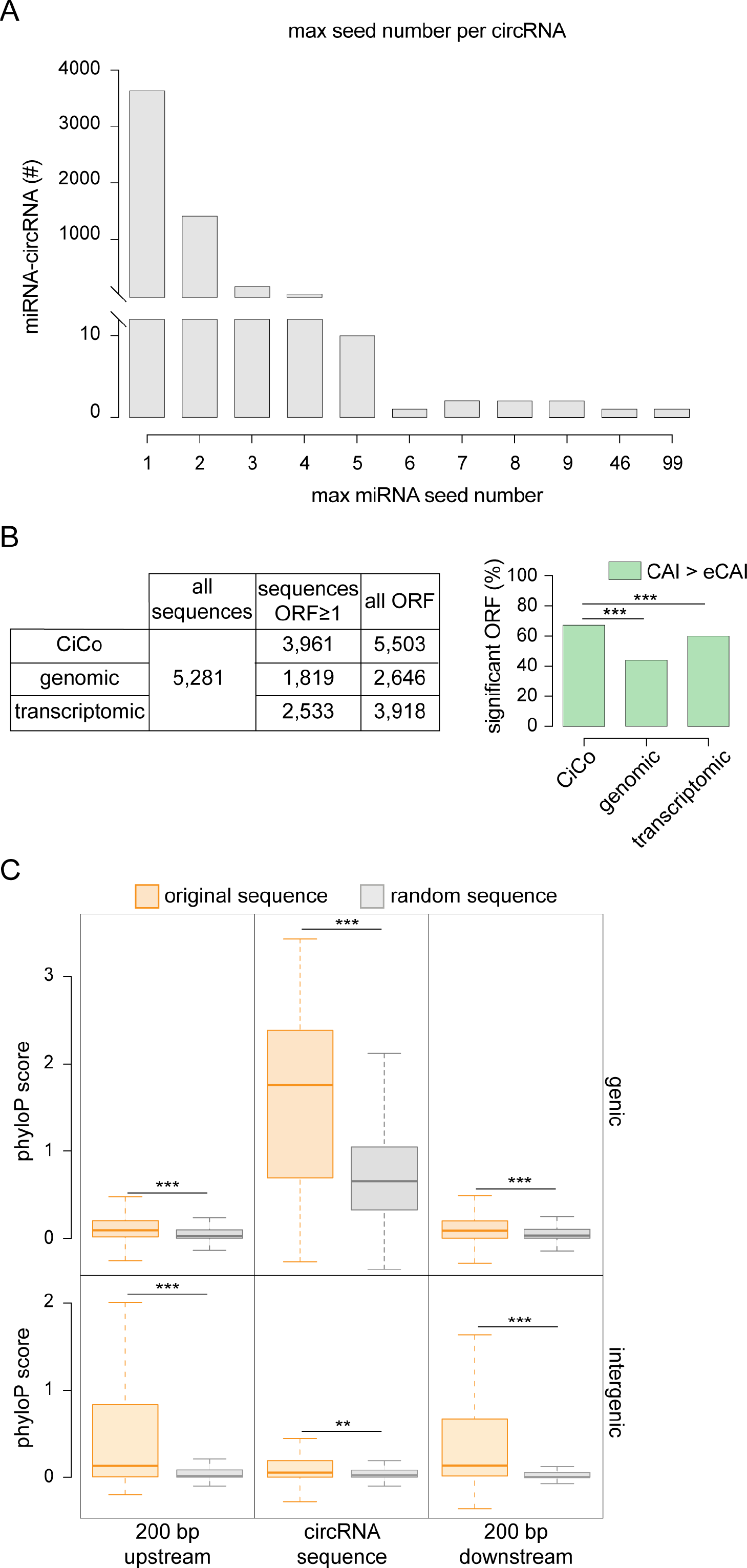
Seed sequences, coding potential and evolutionary conservation of circRNAs. (**A**) Distribution of the maximum number of seed sequences for a miRNA found across CiCo Note that the overwhelming majority of circRNAs only displayed few (1-4) seeds. (**B**) Total ORFs (left) and proportion of those with a CAI value above expected (right) found in CiCo or random control datasets (*** *p*<0.001; chi-squared test). (**B**) Whiskers-box plots showing the distribution of phyloP scores for CiCo sequences (orange) and relative shuffled controls (grey) (** *p*<0.01; *** *p*<0.001; Student’s t-test).

In addition to miRNA sponging, a recent study has suggested that at least one circRNA, circ-ZNF609, is translated into a peptide ^27^. Hence, we searched for ORFs among all circRNAs and found that a remarkably high proportion (nearly 4,000; i.e. >70%) contained at least one putative ORFs with ≥150 nt in length (**Figure 3B**, left). To increase our confidence in this result, we assessed the Codon Adaptation Index (CAI) measuring the relative codon usage as a function of gene expression ^37^. To this end, we used CAIcal ^38^ to compute the index of each predicted ORF within circRNAs and referred this to the expected value calculated from a pool of 500 shuffled sequences reflecting the whole ORF population (eCAI=0.748) that was used as a significance threshold. We found that 68% of all predicted ORFs and belonging to ~50% of all circRNAs had an index higher than expected (**Figure 3B**, right) raising the possibility that these are potentially translated. However, 96% of CiCo’s sequences derive from coding genes implying that this abundance of ORFs might simply reflect their sharing of some, if perhaps not all, the features of coding genes. To assess this, we repeated our analyses (ORF prediction and CAI calculation) but this time considering random sequences generated not only from the entire genome but also from the coding transcriptome as two independent negative controls. Although we could find a comparable number of putative ORFs from CiCo and our two negative control datasets (6,375, 3,520 and 4,742, respectively), only 48% of all random genomic ORFs and 61% of the transcriptomic ones were found to have a CAI value above the eCAI thresholds. Remarkably, when comparing the fraction of significant and non-significant ORFs within CiCo with the two control datasets, we found highly significant differences with both (Chi-squared test, two-tailed pval<0.001) (**Figure 3B**, right) suggesting that the coding potential of CiCo is not only an inherited feature resulting from their origin from coding genes but might potentially account for their function.

We next assessed the evolutionary conservation of CiCo. To this end, we considered not only the sequence of the mature circRNAs but also their flanking regions (200 bp up- and down-stream) as these are important for their biogenesis ^6,20^. Again to obtain an appropriate reference as negative control, we generated for each circRNA and flanking regions a random sequence of the same size and reflecting the features of such circRNA. Specifically, we used any random genomic region for intergenic circRNAs or transcriptome-specific and intronic sequences for genic circRNAs and their flanking regions, respectively. We next computed the average conservation score of all these regions making use of the phyloP score (a per-base value obtained by the alignment of murine genome against 30 other vertebrates, UCSC ^39^) and compared it with the one calculated for the respective random sequences. We found that genic circRNAs were more conserved than their reference random sequences and that despite an average lower conservation score their flanking regions were also significantly more conserved (**Figure 3C**; top). As this again can be influenced by circRNA origin from coding genes, we next assessed the conservation of the subgroup of intergenic circRNAs that, by definition, are not overlapping any annotated gene. Strikingly, both the circRNAs themselves as well as their 200 bp up- and down-stream regions were also significantly more conserved than their random counterparts (**Figure 3C**; bottom). Hence, similarly to their coding potential, the conservation of circRNAs seems not only an inherited feature resulting from their sharing of sequences with coding mRNAs but may actually reflect their function.

### CircRNA biogenesis is independent from alternative splicing

Finally, since nearly all (96%) CiCo transcripts are overlapping genes, we asked if the abundance of circRNAs correlated with that of their respective linear mRNA, which could reveal the mechanisms underlying their biogenesis. In this context, we sought alternative mechanisms by which biogenesis of circRNAs may be controlled. We speculated that this may occur at the level of transcription by the synthesis of different pre-RNAs that ultimately mature into circular, rather than linear, transcripts. Alternatively, a circular or linear RNA may result by splicing of a single pre-RNA precursor. Although not mutually exclusive, the two mechanisms may occur either un-specifically without any regulation at the level of individual transcripts or cell types or, alternatively, only apply to a specific sub-class of genes. Ultimately, distinguishing between these possibilities may help understanding the biological significance of circRNAs.

To this end, we took advantage of the previous poly(A) transcriptome assessment of PP, DP and N reported by our group ^30,33^ and compared the expression levels of each circRNA and its corresponding linear counterpart in each cell type. We found a strong positive correlation between the expression of mRNAs and circRNAs (Pearson’s score ~0.8 in all cell populations) (**Figure 4A**) implying that an increase in a circRNA did not result in a decrease in its linear form. This in turn led us to reject the hypothesis of regulation at the level of transcription of different pre-RNAs. Next, if synthesis of a circular versus linear RNA should be regulated by splicing from a common transcript, then an increase in such circRNA should inversely correlate with the usage of the exon(s) in common with the linear transcript. To assess this, we performed differential exon usage analyses of PP, DP and N (to be described elsewhere) and assessed whether exon(s) used in common by both a circular and linear transcript would display an inverse FC from one cell population to the following (e.g. from PP to DP and from DP to N). Surprisingly, barely 0.5% of all circRNA-exon pairs showed an opposite FC pattern with regard to their linear counterparts while the vast majority shared the same pattern of either up- or down-regulation (**Figure 4B**). In turn, this questioned a mechanistic link between mRNA splicing per se and circRNA biogenesis. To address this, we next selected all genes resulting in the expression of at least 2 linear isoforms resulting from alternative splicing as one specific mechanism of splicing and investigated what proportion of circular RNA was produced among these genes. We found that only 49% of circRNAs were generated from alternatively spliced genes and that among these only ~0.2% shared the very same exon(s) in all three cell populations (**Figure 4C**).

**Figure 4.**
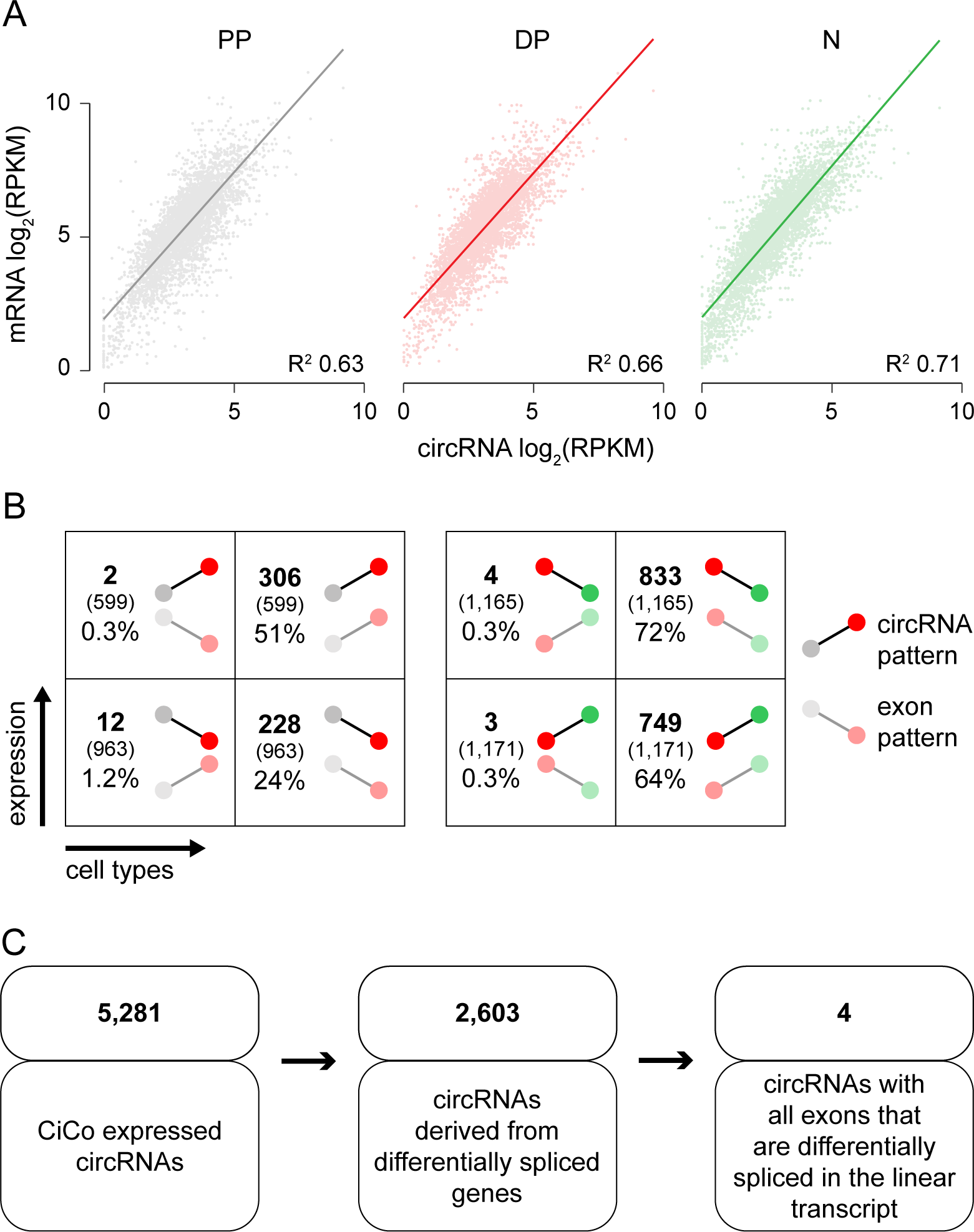
CiCo relation to linear transcripts. (**A**) Scatterplot log_2_(RPKM) of CiCo and mRNA counterparts in cell types (colours as in **Figure 2C**). R^2^ for each regression line are indicated. (**B**) Number and proportion of circRNAs having either an opposite or the same FC pattern as the exon of the linear counterpart (50% i.e.: FC ≥1.5 or ≤0.67; no FDR being applied; colours as in **Figure 2C**). (**C**) Number of expressed circRNAs that result from differentially spliced genes during the neurogenic lineage.

Taken together, our data indicate that the overall expression of circRNAs correlates with that of their linear RNAs but is not a by-product limited to transcripts and exons undergoing alternative splicing and, hence, can be regulated independently by mechanisms that are still unknown.

## DISCUSSION

Here, we provided the arising field of circRNA biology with the first resource describing their sequence and cell-specific expression during mammalian cortical development. This has allowed us to increase by 3-fold the number of circRNAs currently known pointing out their full diversity and specificity in cell populations representing the lineage to neurogenic commitment. More importantly, our work has allowed us to reveal novel features of these elusive transcripts that are important to infer their significance and that could not be deduced based on the previous knowledge of backsplice junctions alone.

We found that genic circRNAs are primarily composed by exons of genes encoding for mRNAs or lncRNAs and that a strong correlation existed in the levels of expression of circular and linear pairs of transcripts. Our data were inconsistent with both the synthesis of circular-specific, pre-RNAs during transcription as well as the involvement of alternative splicing as a mechanism underlying their biogenesis. While more studies are needed to address these aspects, certain features emerging from our study seemed to support the notion that circRNAs may be a generic by-product of splicing while others highlighted their specificity both in terms of biogenesis and putative biological significance. To start with, CiCo transcripts were found to derive only from a specific group of linear transcripts and consistently included certain exons but not others. In addition, both the coding potential and evolutionary conservation of CiCo revealed to be much higher than expected by chance even after accounting for their origin from coding exonic regions.

While it is clear that a significant proportion of circRNAs might nonetheless lack function despite these features, our study provides the field with a new resource for the identification of biologically relevant circRNAs fuelling future studies on the molecular mechanisms underlying their regulation and function.

## MATERIALS AND METHODS

### Animal care, cell sorting and RNA extraction

Animal experiments were approved by the Landesdirektion Sachsen (24-9168.11-1/41 and TVV 39/2015) and carried out in accordance with the relevant guidelines and regulation. Pregnant *Btg2*^RFP^/*Tubb3*^GFP^ double heterozygous mice were anaesthetised with isoflurane and sacrificed through cervical dislocation. Brains from E14.5 embryos were collected and the lateral cortex isolated after removal of the meninges and ganglionic eminences. The Neural Tissue Dissociation Kit with papain (Miltenyi Biotech) was used according to the manufacturer’s protocol to obtain a cell suspension for FAC-sorting or RNA extraction. Cells were resuspended in ice cold PBS and supplied with 10 μl of 7-AAD to assess cell viability (BD Pharmigen) and sorted with a BD Aria III FACS as previously described ^30^. Sorted cells were collected in PBS, centrifuged (2000 rpm, 5 min) and RNA extracted using the Quick RNA Mini Prep (Zymo Research) according to manufacturer’s protocol.

### CircRNA sequencing, annotation and validation

Total RNA was denatured for 3 min at 70°C, RNase R (Epicentre) treated for 1 h at 40°C and supplied with DNase I (Invitrogen) 15 min at room temperature. The reaction was then cleaned with RNA Clean & Concentrator (Zymo Research) and cDNA libraries prepared using the NEB Next Ultra Directional RNA Library Prep Kit without mRNA enrichment. Effectiveness of RNase R treatment was assessed by qRT-PCR of 6 transcripts with the highest free energy per nucleotide in their most stable predicted secondary structure according to RNAfold (-p --noLP --temp=40; primers in **Table S1C**). Samples were sequenced on an Illumina HiSeq 2500 with a read length of 150 bp that were aligned using gsnap ^40^ and mm9 as reference genome. Quality was evaluated by the percentages of mapped and unmapped reads yielding similar values to the only comparable study reported to date ^6^ (**Figure S1B**). To predict putative circRNA, unmapped reads were retrieved and then analysed using the “find_circ” pipeline (version 1) with default parameters ^13^. No filter was applied on the number of reads identifying the circularising junction. For the genomic location, predicted circRNAs were overlapped with mouse ENSEMBL Genes (v67 ^41^) and considered them genic when their start/end base coincided with the start/end of annotated exon(s). circRNA overlapping genes but with other start/end points were grouped together with intergenic and antisense ones (Figure 2A, termed as “other”). Overlap with circBase was done using bedtools ^42^ and intersecting predicted circRNAs with the .bed file for mouse ^35^. Different levels of minimal reciprocal overlap were set within bedtools options to take into account also single-nucleotide overlap between data. For validation, 1μg of total RNA from un-sorted lateral cortex was treated as described previously for sequencing, with one sample digested either with RNase R or water and then converted into cDNA using 200 U of reverse transcriptase and 50 ng of random hexamers according to the SuperScript III (Invitrogen) kit. This cDNA was diluted 1:10 and 1 μl used as PCR template with divergent primers designed with Primer3 ^43^ and spanning over the circularising junction. All primer pairs were tested with the In-Silico PCR tool from UCSC to minimize by-products ^44^ (primers in **Table S1A**).

### Sequence prediction, differential expression and conservation

First, exonic and intronic sequences of previously annotated genic circRNAs were separated. Next, we retrieved the coordinates of these exon(s) from ENSEMBL Genes (v67 ^41^), while intron coordinates were obtained through command line starting from the exonic ones. circRNAs annotated as intergenic, antisense or genic with unusual start/end points were considered as a single exon. Exonic and intronic coordinates were kept separate and featureCounts ^45^ run using as reference files the exon and the intron coordinates to obtain a per-feature read count on which RPKM values were calculated as follows: 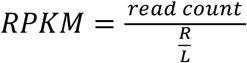, where: *R* = *lobrary size* * 10^−6^ and *L* = *circ length* * 10^−3^. Since validation and cloning was performed on RNA from the total lateral cortex, RPKM values per circRNA were summed and not averaged. Threshold for expression was set as the highest RPKM of predicted, but not detected, introns within 10 validated circRNAs with additional 6 circRNA used to validate the absence of introns from which we redefined the set of expressed features (primers in **Table S1B** and **D**). For the differential expression analysis, we run again featureCounts ^45^ using the set of expressed features as reference and specifying a meta-feature count to automatically sum the reads from different features of the same circRNA. The resulting table was analysed with DESeq2 ^46^ considering fold changes by ≥1.5 or ≤ 0.67 (no FDR applied). Clustering analysis was used to evaluate enriched terms using DAVID ^47^. Finally, circRNA conservation was assessed by computing the phyloP score for the expressed sequence as well as for 200 bp up- and down-stream of each circRNA. To compare our sequences, we generated a shuffled version with the same sizes using bedtools (shuffleBed) and accounting for their genomic location. In particular, for genic circRNAs the shuffled sequences were obtained from a reference file that included exons and/or introns as appropriate and derived from ENSEMBL (v67). In the case of intergenic circRNAs (both sequence and flanking regions), the shuffled sequences were chosen from the entire genome. BEDOPS ^48^ was used to sort regions per genomic location and to split into a per-base coordinates file. BEDOPS was again used to retrieve the per-base phyloP score from the UCSC table containing the conservation score for mouse vs 30 vertebrate genomes (phyloP30wayAll). For shuffled sequences relative to genic circRNAs, we built a custom phyloP score table in which the values relative to either exons were included. We finally reconstructed the score per sequence by averaging the phyloP values for each circRNA. These steps were performed for all files (original and shuffled sequences).

### Prediction of miRNA seeds, ORFs and exon usage correlation

Seed prediction was performed using miRanda ^49^ with default parameters and with input the FASTA file for the circRNA sequences obtained with bedtools (getfasta, with the expressed feature coordinates file as input) and miRBase-downloaded mouse mature miRNA sequences (v22) ^50^. The output generated was then parsed to a more manageable form and the seed category was added according to Bartel ^51^. To count the number of seed for each miRNA present on a circRNA, we first subset the parsed miRanda output by seed category (removed everything that did not have a recognized seed type), alignment score (≥ 150) and by free energy (ΔG≤−19) ^51^ then we counted and reported the number of unique miRNA-circRNA combination and sorted by number of occurrences (top to bottom). We predicted the presence of ORF using the standalone version of NCBI’s ORFfinder ^52^ using the FASTA sequences of the expressed circRNA features as input. We restricted the search by requiring a minimum ORF length of 150nt and setting the start codon as ATG only; we also ignored nested ORFs and searched only on the strand of the sequence itself. We then supplied the resulting sequences to the CAIcal web-server ^38^ together with the Codon Usage for mouse ^53^. Expected CAI value (eCAI) was calculated several times using the Markov method with 95% of confidence and the final value obtained by averaging all the calculated ones. As controls, we generated two shuffled dataset using as reference either the entire genome or the transcriptome only. For this two control datasets, the same ORF prediction and CAI calculation were performed. Differences of our dataset with the two control ones were assessed through Chi-squared test with one degree of freedom. Exon usage data were obtained by poly(A) enrichment and PE sequencing of our 3 cell population (to be reported elsewhere). For the full mRNA we selected the RPKM values for the linear corresponding to the circRNAs. For linear versus circRNA expression, we computed the average RPKM for each cell population and compared the log2(RPKM), plotting their relative abundance and computing an overall Pearson’s correlation score. In case of linear exon versus circRNA, the corresponding read counts were analysed with DESeq2 to compute the FC of the exon shared by circRNAs and linear transcript.

## Data availability

All custom scripts can be obtained upon request. Sequencing data generated during the current study are available at GEO repository (GSE117009).

## ACKNWOLEDGMENTS

We thank the MPI-CBG and CRTD facilities for maintenance of the mouse lines, sequencing and FACS. We are grateful to the members of Silvio Bicciato’s Lab (University of Modena) and Dr. Michael Hiller (MPI-CBG) for suggestions and discussion. We also thank Prof. Katja Nowick and Maria Beatriz Walter Costa for insights into RNA structure prediction. This work was funded by the CRTD, the School of Medicine of the TU-Dresden, the Italian Epigenomics Flagship Project (Epigen), DFG grant DFG CA 893/9-1 and a DIGS-BB fellowship to MD.

## AUTHOR CONTRIBUTIONS

FC and MD conceived the project; MD carried out the bioinformatic analyses supported by ML and AD and all experiments with the help of DC and SM; LHAA provided the differential exon usage data. FC and MD wrote the manuscript. All authors approved the manuscript.

## ADDITIONAL INFORMATION

Supplementary information accompanies this paper. The authors declare no competing interests.

