## Supplementary Information for "Comprehensive Assessment of Sequence and Expression of Circular-RNAs in Progenitor Cell Types of the Developing Mammalian Cortex"

**Figure S1. Related to Figure 1**. (**A**) Validation of RNase R treatment on highly stable expressed transcript. Graphs represent the cycle threshold values (Ct; top) or calculated fold change (FC; bottom) upon qRT-PCR of total RNA from E14.5 mouse cortex, with or without prior RNase R treatment (as indicated). For linear transcripts, regular primers were designed while divergent primers were used for circEzh2. N=3, n=3; bars=SDs (* $p<0.05$; ** $p<0.01$; *** $p<0.001$; Student's t-test). (**B**) Sequencing quality was assessed by comparing the overall statistics (left) with the only other study using RNase R treatment prior to sequencing (Jeck *et al.*, 2013). Total number of reads for each sample are reported (right). (**C**) DNA gel electrophoresis upon PCR amplification of 10 circRNAs predicted to include introns. Divergent primers were designed to include also the intronic sequence. Full-length gel is available in **Figure S2**. (**D**) Expressed circRNAs were identified as genic when overlapping in the sense strand of genes and the start/end base of the circRNA coincided with the start/end base of an annotated exon (top). If genic, but with start/end base falling within and intron or exon we grouped the circular with the antisense and intergenic ones and generically termed them as other (bottom, top to bottom respectively). (**E**) Primer designing strategy. Divergent primers were designed for specific detection of the circular-specific backsplice junction (top), while convergent, regular, primer pairs were used to identify linear, non-circularized, mRNA (bottom).

**Figure S2**. **Full-length gel with PCR products**. Bold circRNA ID represent the original panel for each circRNA, while small-sized ID are either for circRNA for which a second PCR was necessary to obtain enough material for cloning or un-specific product (determined after sequencing of the cloned PCR product). (**A**) Gel shown in **Figure 1**. (**B**) Gel shown in **Figure S1 C**.

**Table S1**. **Primer List**. (**A**) List of divergent primers for all the tested circRNAs in **Figure**

**1C**. (**B**) Primer used for cloning and sequencing in Figure 1B. (**C**) Primers for linear transcript used in RNase R efficacy test, **Figure S1A**. (**D**) List of divergent primers for intron validation, **Figure S1C**.

**File S1**. **Expressed feature**. Tab-delimited file containing exon/intron coordinates, the custom circID assigned and the type of feature expressed (exon or intron) as well as the calculated RPKM in each cell population. **CiCo (Ci**rcular RNA from lateral **Co**rtex**)**. The first 6 columns are in BED format. Additionally, it is reported the overlapped gene name (or Intergenic) and ENSEMBL gene ID, the number of exon and intron included (in case of genic circRNAs; for antisense and for genic with unusual start/end point no exon or intron is reported). In the last two columns are reported the circBase ID (if there is a correspondence) and the minimum level of overlap between the circRNAs. **miRNA seed prediction**. Tab delimited file containing the total number of seed predicted, the relative miRNA, the circRNA id on which the seed was predicted and the length of the circRNA.

**Table S1 (A)**

| CiCo_mm9_circ_ | | Sequence | Expected size |
|---|---|---|---|
| 001074 | FWD | GATGTGAGCCTGACTGAACG | 147 |
| | REV | CTGCTGCTGGTTCTTGCTC | |
| 001268 | FWD | CAGCTTGGCATGTGGAAAT | 146 |
| | REV | CAGAGAAGAGCCCCTGGAC | |
| 001423 | FWD | GCCTTGTTCTTCTGGTGAGC | 154 |
| | REV | CTCCTGAACAGAAGGCTTGAA | |
| 001527 | FWD | CCCTGAAAATGCAGAGAGAA | 142 |
| | REV | TCTTGTTGGGCACAAAGATG | |
| 001902 | FWD | CAAGTGTGACAGGCGCATAA | 143 |
| | REV | CACCTGGTGGTTGATCTTGA | |
| 002029 | FWD | CCAACTACGCAGACCCCTAC | 148 |
| | REV | TCTGCTGAACTGGGGTCTTT | |
| 000203 | FWD | CGCGAGGGACTGACAGAGT | 146 |
| | REV | AGGGTCCGTGCGAAGAAG | |
| 002131 | FWD | GAACCCCAAGGCTCTGCT | 149 |
| | REV | GAACAGTTGGTGACCACGAG | |
| 002132 | FWD | CCTGCTGGACCTCTCAGG | 149 |
| | REV | GGATCTCAGCCAGAGACATGA | |
| 002204 | FWD | GGACAGTGAGGAGCTCAGG | 153 |
| | REV | AGGCTCCGAAGAAAGTGCT | |
| 002231 | FWD | AAGAGCAGGGCATCATCTCT | 149 |
| | REV | TCTTTGGCAAGCTGTGGTC | |
| 002232 | FWD | GGGATGCATCTCTTGATAACTG | 147 |
| | REV | TCCTCCAGATCTCTGTGGAAT | |
| 002259 | FWD | ATCGGAGTTTTGGACCAATC | 145 |
| | REV | TTGCTCGACTCATAGCTGGA | |
| 002390 | FWD | CAACATCTCCTCGGATGTCA | 145 |
| | REV | AGGCCACTTTTGAGTTCTTGTC | |
| 003161 | FWD | CTCACGGGGCTCTGTCAAG | 149 |
| | REV | GGAAGCTCCCATCAGGAAAT | |
| 003578 | FWD | AAACGCTGACATCGAGCTG | 156 |
| | REV | GCGATCGGTTCTTAGCACTC | |
| 003934 | FWD | CAGCAACCACCAGCTCCTAT | 145 |
| | REV | GGCCTGTGATATCATTCTGCT | |
| 003968 | FWD | GGCCCAAACCCTTAAAATTAC | 145 |
| | REV | TTTGCATCTTGCTCCCTCAT | |
| 004187 | FWD | CCTCTCCACCACTGTCAGC | 150 |
| | REV | CATGTCCAGTTCCTCTGAAGAT | |
| 004202 | FWD | GGAAGATGAGGACGAAGATGA | 143 |
| | REV | TGTCACCTGAATTTCGTCTTTC | |

| | | Primer sequence | Expected size |
|---|---|---|---|
| 004254 | FWD | TTACTGTGAAGTTTGCCAACAA | 144 |
| | REV | TGAGGCTCCATGGTGCTAAT | |
| 004860 | FWD | AACACCATCACTCGGCTAAAG | 147 |
| | REV | TGTGCCCAGGATGTTACAGA | |
| 004878 | FWD | ACGTTCCTGCCTGAGCTG | 153 |
| | REV | CACAGGTGCCTTGGTAAGGT | |
| 005387 | FWD | TTGGAGAAAATGCTTCCAGA | 148 |
| | REV | CCAGGATTCAAGCCAGTGTC | |
| 005420 | FWD | TCCCAGGAAGAATTACAAAGG | 171 |
| | REV | TCTGAAGACTTCTGGGTCTGC | |
| 005430 | FWD | GGCTGTCTGACTGGTGGAAC | 147 |
| | REV | CTGTGGACCCAGTGGTGAC | |
| 006088 | FWD | GACTGGCCAGGGGACTTC | 143 |
| | REV | CGGTGGCACCAGAGTGAC | |
| 006218 | FWD | ACCTCTGGGCAGAACAGC | 141 |
| | REV | GTTCCAGATCAAATCCCTTGA | |
| 006222 | FWD | GGGCCATGGTATCTCTGTGT | 147 |
| | REV | CCAGGAATTAGCCAGGATTTG | |
| 000669 | FWD | GCCTGTCTACATCAACATCATC | 151 |
| | REV | GAAGGGAAGGACCAGGTAGC | |
| 000859 | FWD | ATGAACCAGCTTTCCCTCCT | 145 |
| | REV | TAGCTGACGAGCCCTCTCTC | |
| cdr1as | FWD | CTCCAGTGTATCGGCGTTTT | 153 |
| | REV | TCACGATTGTCTGGAAGACCT | |
| 000202 | FWD | GCCATTCAGGCTCATCAATA | 156 |
| | REV | GTCTCGGTCATTCCACGACT | |
| 000239 | FWD | GCTACCCCAGCTCCAACAT | 147 |
| | REV | GCTTAAGAGGGCTGTGCTGT | |
| 005390 | FWD | ACAGCAGACAGCTCCAATCA | 150 |
| | REV | TTGGAAAGATGGGTGTTGGT | |
| 006087 | FWD | CCCTGAACGACTGTATGCAC | 149 |
| | REV | GGCACGGAAATCCAAGCTAT | |
| 002232 | FWD | TGTGGCCTGTGTATGAAGGA | 151 |
| | REV | CTCCAGAGGCCAGTAAGTTCC | |
| 003872 | FWD | TCCTTCTTCAGCAAACACAACA | 149 |
| | REV | TGAGATTTCGAGCTTGTTTGG | |
| 000720 | FWD | GCTTCAACTGGAATGGCAAG | 154 |
| | REV | TCTGGGCCATGTCTGAATAA | |
| 000721 | FWD | TGTGCCTTCATTGCACATT | 155 |
| | REV | TCTGGGCCATGTCTGAATAA | |
| circEzh2 | FWD | TTACACGCTTCCGCCAAC | 131 |
| | REV | AAGCAGCGGAGGATACAGC | |

**(B)**

| CiCo_mm9_circ_ | | Primer sequence for cloning | Expected size |
|---|---|---|---|
| 000203 | FWD | ACGTCTCGAGAGGACCCTGCTTTCTCTGCTGTGATTC | 37,144 (701) |
| | REV | ACGTGTCGACACCGTGCGAAGAAGGAAGCGGCCACTG | |
| 000720 | FWD | ACTGCTCGAGATACTCAGGTTGATGGAGTCAGGGA | 32,985 (961) |
| | REV | ACGTTCTAGACTTTGGCTGAGCAAGTTTACAGTTT | |
| 000721 | FWD | ACTGCTCGAGATACTCAGGTTGATGGAGTCAGGGA | 27,052 (754) |
| | REV | CAGTGTCGACCTTGATAAGCAGAACTTAGCCATTT | |
| 003872 | FWD | ACTGCTCGAGCGCCATCATATAGGAGATCGTAGCC | 5,307 (611) |
| | REV | CAGTGTCGACCAGTTTGTATACAGACTCGTGATGG | |
| 004202 | FWD | ACTGCTCGAGAGGTGACAGAGTTAGTCCTCGATAATT | 4,131 (603) |
| | REV | CAGTGTCGACACCTGAATTTCGTCTTTCATTAAGTAT | |
| 004583 | FWD | ACGTCTCGAGAGCCAGGTCGCACTGCCTGCGTCACTA | 199 |
| | REV | ACGTGTCGACACGAGGGACATGTTCCTTATCCTTTCG | |
| 006087 | FWD | ACGTCTCGAGAGGCCTCAGTATGGTGGCAAGTACTGT | 6,529 (673) |
| | REV | ACGTGTCGACACTTCAAAGACCAGCGTCTCATTCGTG | |
| 006088 | FWD | ACGTCTCGAGAGGACATTTGCAAGTCACTCTGGTGCC | 17,979 (918) |
| | REV | ACGTGTCGACACTTCAAAGACCAGCGTCTCATTCGTG | |
| 006222 | FWD | ACGTCTCGAGAGGCAGGCTAACGAAGAATATCAAATC | 33,824 (490) |
| | REV | ACGTGTCGACACCACTTGTCCATTGTGTGGGTTCTTA | |
| 002131 | FWD | ACGTCTCGAGAGGCCCGCACCTCGTGGTCACCAACTG | 2,606 (257) |
| | REV | ACGTGTCGACACCGTGCGGTTTTTTGACTGCAGCTCA | |
| 004187 | FWD | ACGTCTCGAGAGAAAATCTTCAGAGGAACTGGACATG | 3,711 (331) |
| | REV | ACGTGTCGACACCTTTCTCTTCCTCGGAATGGGCTCA | |
| 005387 | FWD | ACGTACGCGTAGGAGGAAGAATATGGAAAAGACAATG | 17,031 (3,784) |
| | REV | ACGTTCTAGAACCTGAAATTGGAAAGATGGGTGTTGG | |
| 005390 | FWD | ACGTGAATTCAGAAAATGCTTCCAGACTGCTCACTTT | 8,903 (296) |
| | REV | ACGTGTCGACACAGGCTACGATATTGACGTATCTGTT | |
| 005431 | FWD | ACGTCTCGAGAGAACTTTGTGCGTCACCACTGGGTCC | 87,653 (1,409) |
| | REV | ACGTGTCGACACTCATTGAGCAAAGGCATCGAGGTTC | |
| 006631 | FWD | ACGTCTCGAGAGCCATGGAAACAAAGAAGTATTCTCG | 6,546 (375) |
| | REV | ACGTGTCGACACTTGTCATTGACAAAGGAATACATCA | |

**(C)**

| Transcript ID | | Primer sequence | Expected size |
|---|---|---|---|
| circEzh2 | FWD | TTACACGCTTCCGCCAAC | 131 |
| | REV | AAGCAGCGGAGGATACAGC | |
| $Ezh2^{LIN}$ | FWD | GCGGGACTAGGGAGTGTTC | 154 |
| | REV | TGTAAAACAGTTTCGTCTTCCA | |
| GAPDH | FWD | AGGTCGGTGTGAACGGATT | 147 |
| | REV | CGTGAGTGGAGTCATACTGGA | |

| Transcript ID | | Primer sequence | Expected size |
|---|---|---|---|
| ENSMUST00000045942 Emx1 | FWD | CTCACTCTTTCTTCAGCGCC | 170 |
| | REV | CGAGAAGGCTGTGCGAATC | |
| ENSMUST00000056403 H1fx | FWD | CGCACAAGAGCAAGAAGGC | 157 |
| | REV | GGGCGGATAGGGATAGAGAC | |
| ENSMUST00000038537 Wtip | FWD | GATGATTCTGCAGGCCCTTG | 164 |
| | REV | CACAGGAGGCACATTTTGGT | |
| ENSMUST00000052281 A19Rik | FWD | TATGAGCGTCGGACCTCTTC | 163 |
| | REV | TCGGGTGCTTGAAGATCACT | |
| ENSMUST00000154977 Ccdc120 | FWD | TAGGGAGCAGGCGAGGAG | 179 |
| | REV | GCTGGGCAGACTTACAACAC | |
| ENSMUST00000012161 Scarf2 | FWD | GGGGATGAGTGTGGGATAGC | 150 |
| | REV | GTCAGGGCCCCAGAACTG | |

**(D)**

| CiCo_mm9_circ_ | | Primer sequence for cloning | Expected size |
|---|---|---|---|
| 002805 | FWD | ACTGGCTAGCAAGGCAAAATTGGTGTATGAAGAAG | 720 (167) |
| | REV | CAGTCTCGAGTTTTCGATTTTTCTGATGCTGTAAC | |
| 004996 | FWD | ACTGGCTAGCACGGACTCAGACATTGAACAAGGAG | 496 (186) |
| | REV | CAGTCTCGAGTCATCGTCCCATTCAAAGCCTCCGC | |
| 004228 | FWD | ACTGGCTAGCAGGAGCTCTGGTGGCCTGCTGCATA | 940 (547) |
| | REV | CAGTCTCGAGAGGTGAAGCGGGCCTGAAGGTAGAG | |
| 004227 | FWD | ACTGGCTAGCCATGTTCACTATGCTGTATCTGGTG | 883 (490) |
| | REV | CAGTCTCGAGGGCGTGGCCCGAGAAGAAGGACTTC | |
| 003247 | FWD | ACTGGCTAGCCCAGCTGCAGACTTTCTCAGAGGAG | 791 (126) |
| | REV | CAGTCTCGAGGGATTGACAGCAGCCCCCGGTGCTC | |
| 003141 | FWD | ACTGGCTAGCCCTTGGCGAGTGGCAGCCCCTTGAG | 563 (461) |
| | REV | CAGTCTCGAGGGTTCCACCAGTTGGCTGAGGCACC | |

### Dori *et al.* Figure S1

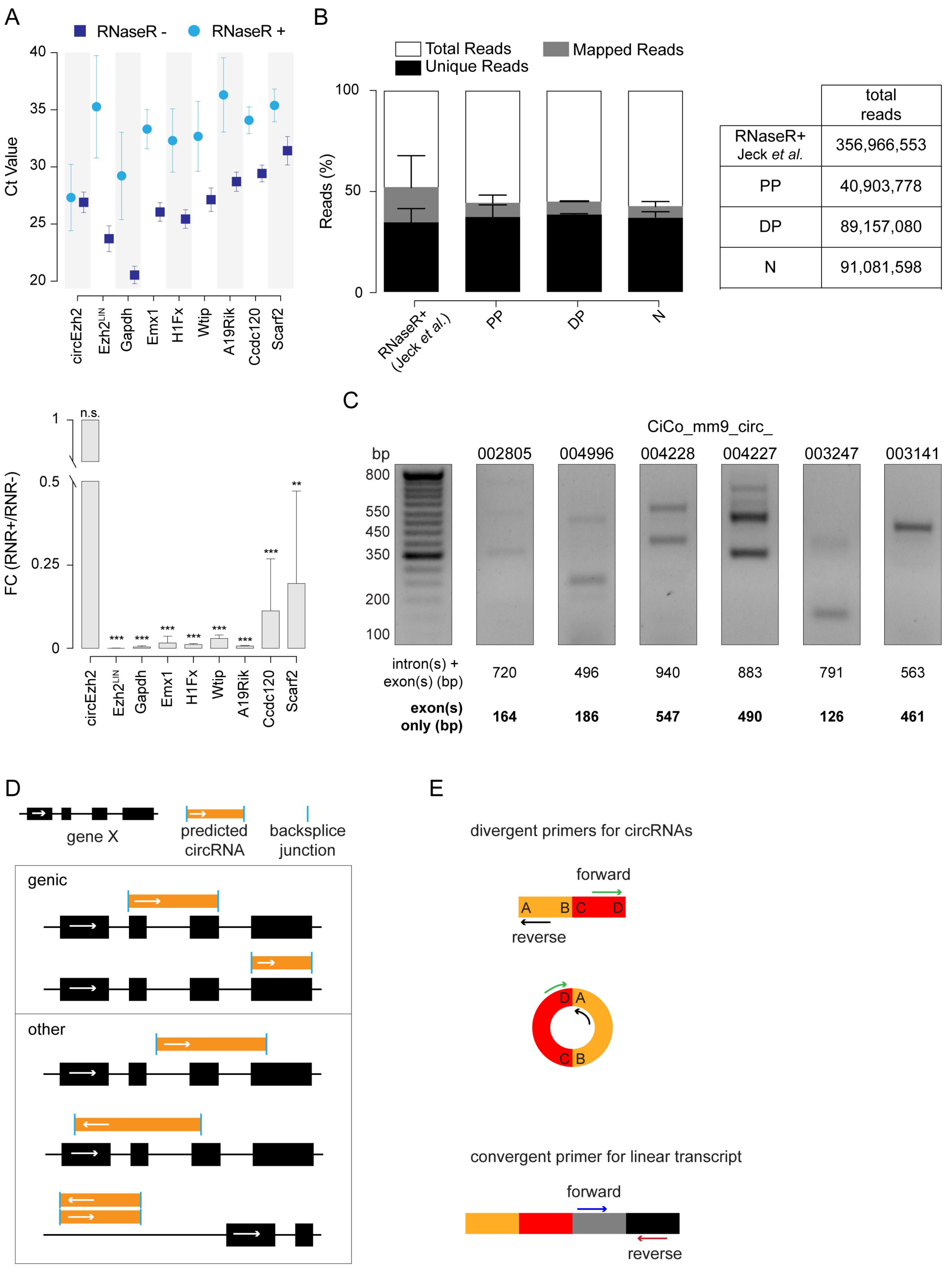


| | total reads |
|---|---|
| RNaseR+ Jeck *et al.* | 356,966,553 |
| PP | 40,903,778 |
| DP | 89,157,080 |
| N | 91,081,598 |

### Dori *et al.* Figure S2

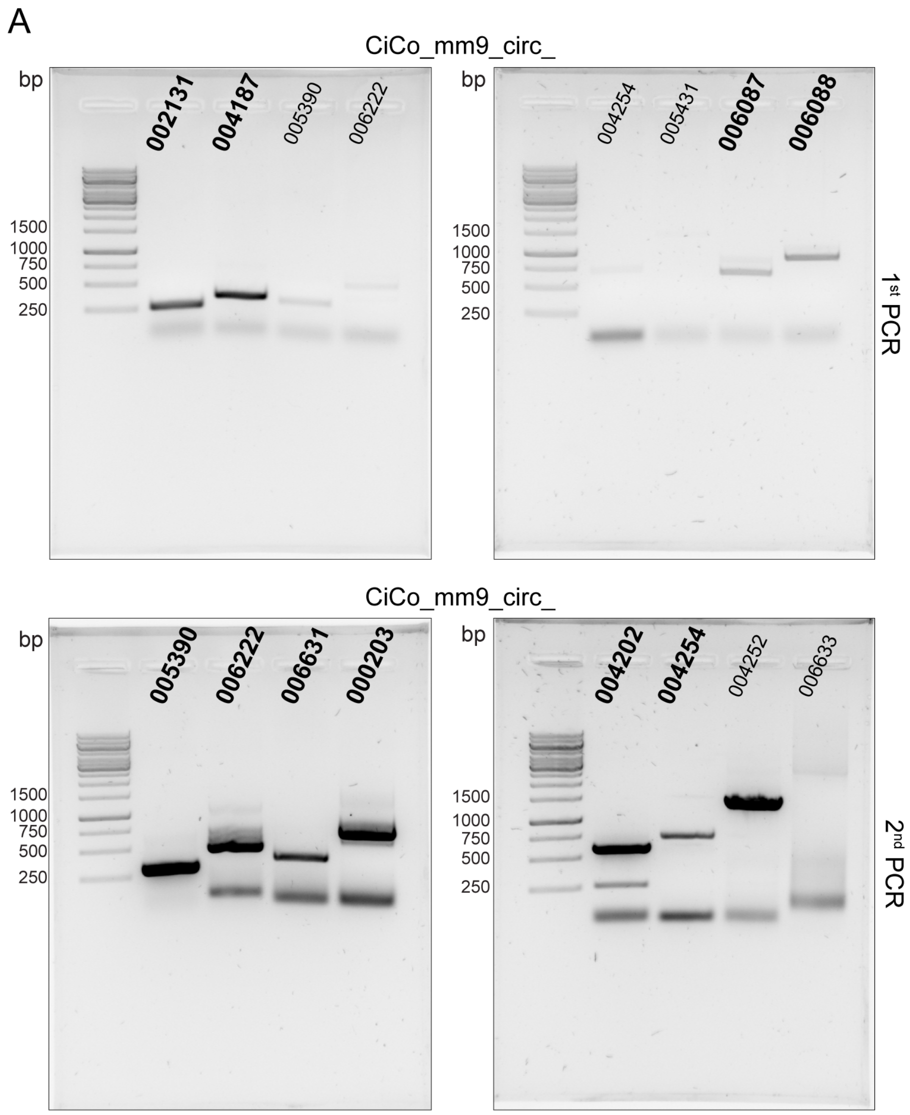


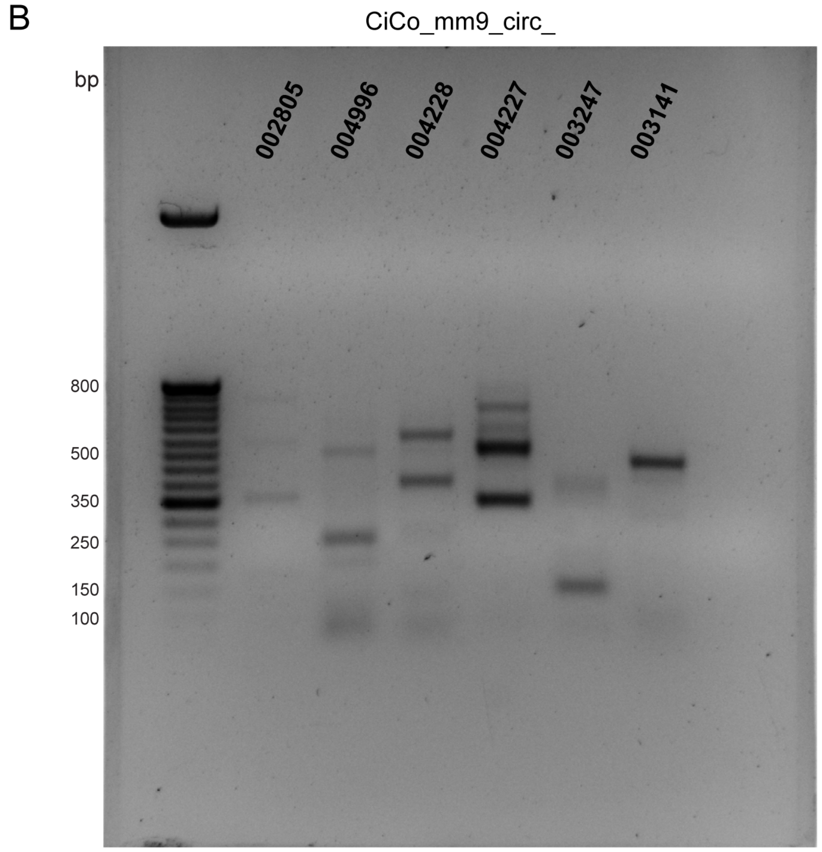
